# Genomic variation of Dutch traditional cattle breeds captured in a Dutch cattle pangenome

**DOI:** 10.64898/2026.11.23.768688

**Authors:** Rensco A.H. Hogers, Mirte Bosse, Richard P.M.A. Crooijmans, Christopher D. Barratt, Bert Dibbits, Marta Gòdia, Annemieke P. Rattink, Aniek C. Bouwman

## Abstract

Traditional livestock breeds are reservoirs of genetic variation lost elsewhere by selection for productivity, yet are rarely represented in reference genome resources. Traditional cattle (Bos taurus) in the Netherlands are severely endangered, with some populations having declined by up to 95 % between 2020 and 2026. We generated 12 assemblies from seven traditional Dutch breeds and combined them with a Holstein Friesian assembly and the Hereford cattle reference to build the first Dutch cattle pangenome (2 881 293 997 bp), a graph containing 193.4 Mb of sequence absent from the international reference. More than 80 % of this sequence was breed private, and this fraction rose with variant size to 99 % for segments of at least one kilobase. Genotyping 54 short-read sequenced animals of these Dutch breeds against the graph recovered 55 865 structural variants and resolved four genetic clusters. The Groningen White Headed was the most differentiated breed while carrying the lowest nucleotide diversity, whereas the recently derived Improved Red and Deep Red remained close to their Meuse-Rhine-Yssel progenitor. Holstein Friesian formed no cluster of its own and contributed the least amount of novel sequence, a reflection of its historical use in upgrading traditional breeds. Ten structural variants located within genes were fixed within a single breed and ranked among the most breed-differentiated of all 55 865 SVs by *V*_*ST*_, including a 174 bp deletion in KIT in the Dutch Belted. Pangenomes recover breed-defining variation that linear references cannot represent, supporting conservation and selection in traditional cattle.

## Introduction

Short-read sequencing remains the workhorse of population-scale genotyping and resolves single nucleotide polymorphisms (SNPs) and small indels accurately and affordably, but it is limited by its read length troubling the detection of structural variants (SVs), particularly large insertions and variation within repetitive regions (Ebler et al., 2022).Long-read technologies, such as Oxford Nanopore and PacBio sequencing, are able to completely span such regions, allowing structural variation to be resolved directly from assembled haplotypes rather than inferred from alignment signatures. Linear assemblies, however, are constrained by the inclusion of only a single haplotype; representing the variation carried across a population requires a framework in which multiple assemblies can be combined, which can be achieved using a pangenome.

Unlike single linear reference assemblies, which inevitably miss variation present in the broader population, pangenomes integrate multiple genome assemblies to represent the full spectrum of genetic diversity within a species (Paten et al., 2017).This is particularly consequential for uncovering the non-core genome, sequence that is absent from the reference but present in some individuals, and which harbours rare variants and structural variation that are invisible to linear reference-based approaches. In cattle, an early multi-assembly bovine pangenome revealed more than 70 Mb of sequence absent from the *Bos taurus* linear reference genome, including genes involved in immune response (Crysnanto et al., 2021). Extending this to global diversity, a pangenome incorporating African breed assemblies added 116 Mb of sequence, improved read mapping rates, and reduced allelic bias (Talenti et al., 2022). Recently, a large-scale pangenome integrating whole-genome sequencing from over 2400 individuals across 82 breeds highlighted the critical contribution of structural variants to cattle adaptation and phenotypic diversity (Dai et al., 2025). However, despite this progress, traditional breeds across Europe remain largely underrepresented in existing pangenomes. A recent effort incorporating French cattle breeds has begun to address this gap (Sorin et al., 2026), but for many regions, including the Netherlands, dedicated pangenome resources are entirely absent.

The breeds least represented in current genomic resources are also, in many cases, those whose populations are declining most steeply. The prioritisation of high-production traits in modern farming has marginalised traditional breeds and reduced their population sizes (Medugorac et al., 2009; Biscarini et al., 2015). Over the last century, the replacement of many local, dual-purpose breeds with a few specialised dairy breeds has affected the genetic diversity of cattle (Hulsegge et al., 2022).This trend is particularly evident in the Netherlands, where in 2020 six out of seven traditional cattle breeds were marked as endangered, with only the Meuse-Rhine-Yssel classified as not at risk (Centrum voor Genetische Bronnen, 2020).By August 2026 traditional Dutch cattle population sizes had declined by as much as 95 % compared to 2020, with the Improved Red falling from 654 to 30 female breeding animals and now classified as critical (Centrum voor Genetische Bronnen, 2020, 2026).The Deep Red and Dutch Friesian Red retain only 249 and 316 female breeding animals respectively, and with the recent decline it is possible that these breeds could be lost in the near future (Centrum voor Genetische Bronnen, 2026).Therefore, it is of the utmost importance to preserve the genetic diversity within these rare traditional breeds, and to characterise what makes them genomically distinct before that diversity is lost. The traditional Dutch breeds are not independent lineages but a closely related group, sharing much of their ancestry and shaped by the same regional breeding history (van Breukelen et al., 2019; Schmidtmann et al., 2021). Because these breeds share so much of their ancestry, a single graph spanning the group can capture that shared backbone once while retaining the sequence that distinguishes each breed, making a pangenome a natural representation for a set of such related traditional breeds.

To produce such a graph we generated *de novo* genome assemblies for the seven traditional Dutch cattle breeds and combined these with existing assemblies of the Holstein Friesian and Hereford into the first Dutch cattle pangenome. We first examined what this graph captures that the linear cattle reference did not, quantifying the non-reference sequence it contains and examining specific loci at which a single-haplotype reference fails to represent or recover a variant. We then characterised the genetic diversity present within and between the traditional breeds, using Holstein Friesian as a specialised commercial outgroup against the distinctiveness of the traditional breeds. Finally, we identified structural variants private to individual traditional breeds and absent from the current reference genome, and considered their potential contribution to traits that define these breeds.

## Materials and Methods

### Sampling and Sequencing

This study included 68 animals, representing seven traditional Dutch cattle breeds and a Holstein Friesian outgroup. To facilitate genome assembly while maintaining a broader population context, we utilised a tiered sequencing strategy: 14 animals with both ONT long-read (LR) and short-read (SR) sequencing, complemented with 54 animals sequenced using SR technology only (Table 1).

**Table 1.**
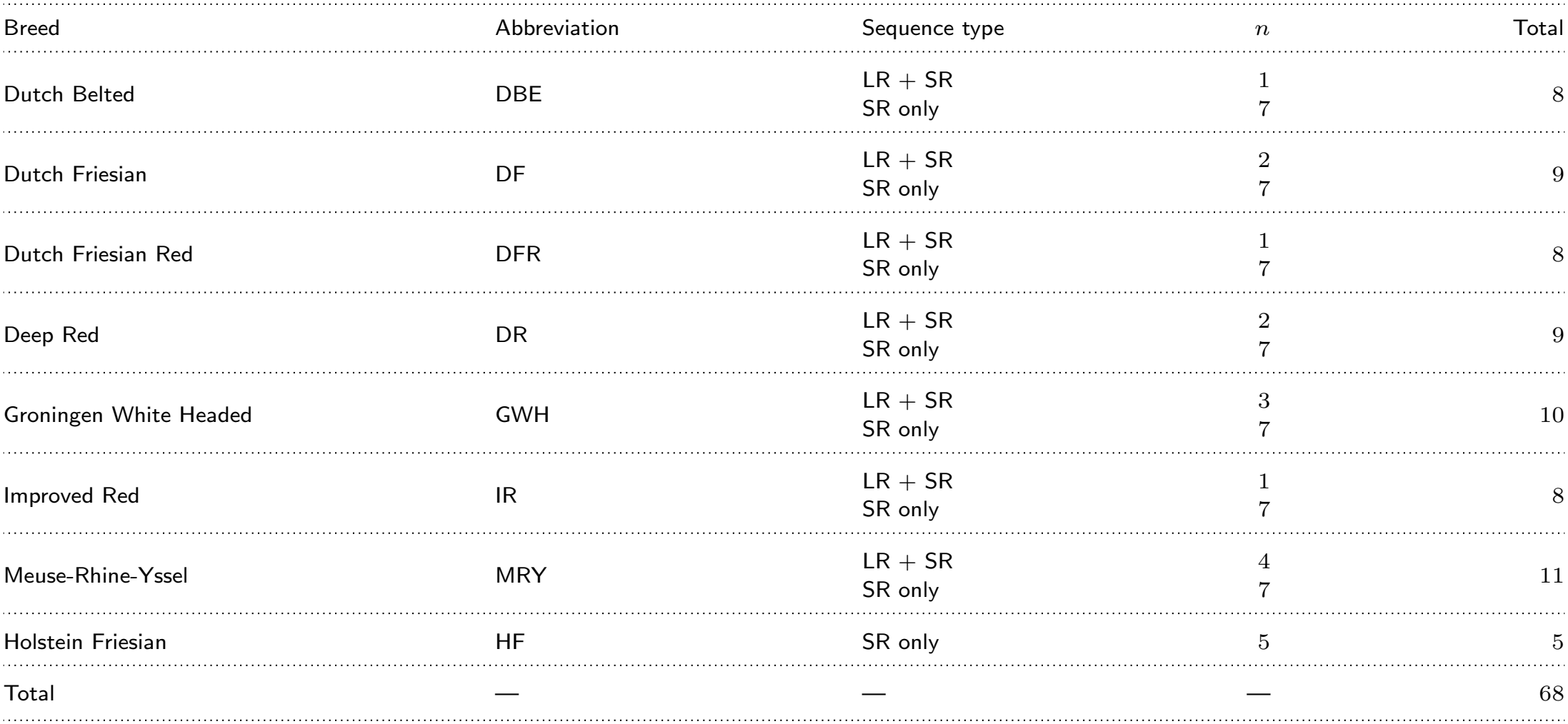
Summary of animal samples per breed. Sequencing technologies include long-read (LR) and short-read (SR).

| Breed | Abbreviation | Sequence type | *n* | Total |
|---|---|---|---|---|
| Dutch Belted | DBE | LR + SR | 1 | 8 |
|  |  | SR only | 7 |  |
| Dutch Friesian | DF | LR + SR | 2 | 9 |
|  |  | SR only | 7 |  |
| Dutch Friesian Red | DFR | LR + SR | 1 | 8 |
|  |  | SR only | 7 |  |
| Deep Red | DR | LR + SR | 2 | 9 |
|  |  | SR only | 7 |  |
| Groningen White Headed | GWH | LR + SR | 3 | 10 |
|  |  | SR only | 7 |  |
| Improved Red | IR | LR + SR | 1 | 8 |
|  |  | SR only | 7 |  |
| Meuse-Rhine-Yssel | MRY | LR + SR | 4 | 11 |
|  |  | SR only | 7 |  |
| Holstein Friesian | HF | SR only | 5 | 5 |
| Total | — | — | — | 68 |

Samples were sourced from two data generation initiatives: ReDiverse (Calus et al., 2017)and OPTIBOV (https://www.optibov.org/). ReDiverse SR samples were sperm-derived, with DNA isolated from semen obtained from the Centre for Genetic Resources, the Netherlands, using the Illumina DNA PCR-Free Prep kit (Illumina Inc., USA). DNA libraries were prepared using the Illumina HiSeq platform (Illumina Inc., USA) at 10*×* coverage with paired-end 150 bp reads. For OPTIBOV animals undergoing SR sequencing, DNA was extracted from blood using the NucleoSpin® Tissue Kit (BIOKÉ) with Proteinase K. OPTIBOV SR libraries were prepared using the Illumina DNA PCR-Free Prep kit (Illumina Inc., USA) and paired-end sequenced (150 bp) on the Illumina NovaSeq6000 (Illumina Inc., USA).

For OPTIBOV animals undergoing LR sequencing blood was collected following the standardised protocols of the OPTIBOV project and DNA was extracted using the GENTRA Blood kit (Qiagen N.V.). For the ReDiverse samples that were subjected to LR one or two frozen straws of semen per cattle were used for DNA extraction.

First, sperm cells were thawed at 37 °C for 30 sec and assessed at the Computer-Assisted Sperm Analysis. Then, semen was washed with Phosphate-Buffered Saline (PBS) and centrifuged at 1500 g for 5 min at 20 °C. DNA was extracted following the protocol as in Gòdia et al. 2023.

For long-read sequencing (*N* = 5 blood, *N* = 9 sperm) DNA quality and quantity were evaluated using 0.6 % (w/v) agarose gel electrophoresis and a Qubit dsDNA Broad Range kit (Thermo Fisher Scientific, USA). DNA from blood tissue was also subjected to shearing using g-TUBE (Covaris, USA) centrifuged at 1200 *×g*, then size-selected using Short Read Eliminator XS (Pacific Biosciences of California Inc., USA) to remove fragments under 10 kb. DNA from sperm was subjected to Short Read Eliminator XS (Pacific Biosciences of California Inc., USA) directly. For both tissue types, library prep were prepared using the SQK-LSK110 kit (Oxford Nanopore Technologies plc, GB) according to the company’s guidelines, except for DNA repair and end-prep step where the incubation step was 10 min at 37 °C and the AMPure XP beads were incubated for 15 min at room temperature. Libraries were sequenced on a PromethION using an R9.4.1 flow cell with two to three loadings per sample (Oxford Nanopore Technologies plc, GB).

#### *De novo* assemblies

*De novo* genome assemblies were generated for 14 animals by using a hybrid assembly-polishing approach. Raw ONT data were converted from FAST5 to POD5 format (v0.3.23) (Oxford Nanopore Technologies, 2024) and basecalled by using Dorado (v0.8.3) with the dna r9.4.1 e8 sup@v3.6model and with adapter and primer trimming enabled. The resulting reads were filtered with Filtlong (v0.2.1) (Wick, 2017) using `--min length` 1000 `--min mean q` 85. Contigs were assembled by using Flye (v2.9.5) (Kolmogorov et al., 2019) and polished for two rounds with NextPolish (v1.4.1;Hu et al. 2020) by using both the long-reads and trimmed Illumina short-reads (Trimmomatic v0.39 `ILLUMINACLIP:TruSeq3-PE.fa:2:30:10 LEADING:20 TRAILING:20 SLIDINGWINDOW:6:28 MINLEN:50;` usadel-lab n.d.). Polished contigs were then scaffolded against the ARS-UCD2.0 reference assembly (GenBank Bioproject PRJNA391427) by using RagTag scaffold (v2.1.0) with default parameters (Alonge et al., 2022), which orders and orients contigs without breaking them and joins adjacent contigs with 100 bp N-gaps. Contig headers were standardised to the PanSN specification with Fastix (Garrison, n.d.). Repeats were identified and soft-masked using the Dfam TE Tools container (v1.90) (Dfam-consortium, n.d.), generating a breed-specific library with RepeatModeler (v2.0.6) (Flynn et al., 2020) followed by RepeatMasker (v4.1.7-p1) (Smit et al., 2013–2015). Assembly completeness and continuity were assessed with BUSCO (v5.8.2) (Manni et al., 2021) using the *cetartiodactyla odb10* data-sets, and QUAST (v5.3.0) (Mikheenko et al., 2018). Read quality was monitored with pycoQC (v2.5.2) (Leger and Leonardi, 2019) and FastQC (v0.12.1) (Andrews, n.d.), and aggregated with MultiQC (v1.27.1) (Ewels et al., 2016). The workflow was implemented in Snakemake (v8.29.2) (Mölder et al., 2021)and is available at Codeberg.

### Pangenome construction

The pangenome graph was constructed with Minigraph-Cactus (v3.1.4) (Hickey et al., 2024)from 12 *de novo* assemblies of traditional Dutch cattle, a publicly available Holstein Friesian assembly (PRJNA668863),and ARS-UCD2.0 (Hereford) as the reference path, giving 14 genomes of which 13 contribute non-reference paths. Assemblies were selected based on N50, total length, and BUSCO completeness, retaining at least one per breed; contigs that did not scaffold to a chromosome were excluded to reduce graph complexity. All subsequent analyses used the default clipped graph.

### Characterisation of non-reference sequence

To quantify sequence carried by traditional Dutch breeds but absent from the linear reference, non-reference content was measured at the level of graph nodes. Nodes not traversed by the ARS-UCD2.0 reference walk were extracted from the clipped GFA, and each was assigned the set of breeds whose assembly walks (GFA W-lines) traversed it. Nodes were then classified as breed-private (traversed by assemblies of a single breed only), multi-breed shared (traversed by assemblies of more than one but not all breeds), or pan-breed (traversed by at least one assembly of every breed), and stratified by node length into five classes: *≤*5 bp, 6 bp to 49 bp, 50 bp to 199 bp, 200 bp to 999 bp, and *≥*1 kb. Because Meuse-Rhine-Yssel and Groningen White Headed contribute four and three assemblies respectively while all other breeds contribute one, breed-private totals were recomputed after downsampling to a single assembly per breed. As a complementary, allele-level view, every ALT allele of the Minigraph-Cactus VCF was classified as a SNP, insertion, deletion, or multi-nucleotide polymorphism from the REF/ALT length difference, with symbolic alleles sized from INFO/SVLEN, and assigned to the same length strata and sharing categories.

To characterise the biological composition of the non-reference sequence, nodes of at least 200 bp were extracted in FASTA format and queried with BLASTN (v2.17.0) (Camacho et al., 2009) against the ARS-UCD2.0 bovine RNA reference set and the Dfam repeat library (Dfam-consortium, n.d.),using an E-value threshold of 1 *×* 10^*−*10^ and a minimum sequence identity of 90 % (RNA) and 80 % (repeats), and alignment over at least half the node length. The highest-scoring hit per node was retained by bit score, and nodes were classified as matching transcript sequence only, repeat sequence only, both, or neither.

### Short-read genotyping against the Dutch cattle pangenome

Short-reads of all 54 genotyped animals were adapter- and quality-trimmed with Trimmomatic (v0.39) using the parameters described in the *de novo* assemblies subsection. The VCF file generated by Minigraph-Cactus was reduced to top-level bubbles with vcfbub (v0.1.2) (Garrison, 2022), discarding bubbles exceeding 100 kb, and further modified to filter out ambiguous alleles with BCFtools view and haploid genotypes were converted to diploid using a custom awk script (Aho et al., 2023).Seven short-read individuals per traditional Dutch breed and five Holstein Friesian individuals (54 animals in total) were genotyped against this VCF with PanGenie (v4.2.1) (Ebler et al., 2022)using the default 31-mers.

### Validation of the pangenome

To test the effectiveness of the Dutch cattle pangenome for investigating structural variants (SVs), we chose a known Copy Number Variant (CNV) found in the Dutch Belted, which underlies the belted phenotype in cattle (Awasthi Mishra et al., 2017; Rothammer et al., 2018). The known location of the CNV was investigated in the genotyped results to check which individuals of the included breeds were annotated with this CNV. Furthermore, the raw VCF output from Minigraph-Cactus was investigated to see if the Dutch Belted assembly was incorporated correctly into the pangenome graph.

To test the effectiveness of the Dutch cattle pangenome concerning SNPs, known Groningen White Headed breed specific SNPs were used. SNPs described by Gonzalez-Prendes et al. (2022), found on BTA20 in the genomic region of the *ZSWIM6, NDUFAF2, PDE4D* and *RAB3C* genes, were queried in a *±*25 bp window. Complex alleles were decomposed with BCFtools norm -a (Danecek et al., 2021), so that markers represented as multi-nucleotide or complex alleles were recovered rather than scored as absent. As a comparison the same short-read cohort was independently genotyped using the linear reference. Trimmed reads were aligned to ARS-UCD2.0 with BWA-MEM2 (v2.3) (Vasimuddin et al., 2019), duplicates were marked with SAMtools (v1.24) (Danecek et al., 2021), and all 54 animals were called jointly across the BTA20 interval containing the nine markers with BCFtools `mpileup and call -m`, retaining reads and bases with a minimum mapping and base quality of 20. Calls were left-aligned and decomposed with BCFtools `norm -m -any -a`, matching the treatment of the pangenome VCF, and genotypes were compared per sample at each marker.

### Genetic diversity analysis

To characterise genetic diversity within and between traditional Dutch cattle breeds and compare these to commercial breeds, genetic variation was assessed by using the genotyping results derived from the pangenome graph. Variant files were merged, normalised, and annotated with INFO tags by using BCFtools. After merging, monomorphic variants and variants located on non-autosomal chromosomes were removed from the sample set for all subsequent analyses.

Within-breed genetic diversity was quantified using observed heterozygosity and nucleotide diversity (*π*) in 100 kb windows with VCFtools (v0.1.17) (Danecek et al., 2011). Observed heterozygosity was derived per individual from the `--het` output as the number of heterozygous sites (`N SITES` minus O`(HOM)`) divided by the autosomal length (2489.4 Mb) and is reported as heterozygous sites per Mb; *π* is the mean of the 100 kb window estimates. Both were estimated from biallelic autosomal SNPs that were polymorphic across the 54 shortread animals. No additional filters on read depth, mapping quality, genotype quality, minor allele frequency or missingness were applied, because genotypes were produced by PanGenie rather than read-level variant calling. Breed-private non-reference nodes (Table S1) were not used directly in these estimates; they contributed only as alleles of the graph bubbles in which they reside, which PanGenie genotyped in every animal regardless of the breed of the assembly that contributed them. Runs of homozygosity were called on the unpruned biallelic SNP set (no LD pruning or MAF filtering) with the hidden Markov model implemented in BCFtools roh, with allele frequencies estimated from the 54 genotyped animals using BCFtools +fill-tags. Genotypes were used directly `(-G30)`, and segments with a quality score below ten were discarded. No minimum segment length was imposed; *F*_*ROH*_ was calculated as total ROH length divided by the autosomal genome length (2.49 Gb) for all segments and for segments ≥100 kb, 1 Mb and 5 Mb.

To characterise breed-level genetic distinctiveness, breed-specific variants were defined as variants present in at least two individuals of the focal breed while being completely absent from all sampled individuals of the remaining breeds. Allele counts were computed per breed group using BCFtools `+fill-tags`, focal-breed alleles were intersected with alleles absent from all other breeds using BCFtools isec matching on exact position, reference, and alternative allele.

Genetic differentiation between breeds was quantified by using the Weir and Cockerham weighted fixation index (*F*_*ST*_) estimator for biallelic sites in VCFtools (Danecek et al., 2011; Weir and Cockerham, 1984). The weighted (ratio-of-averages) estimator were used throughout, which is less biased than the per-site average when sam- ple sizes are small and low-frequency variants are abundant (Bhatia et al., 2013). Genome-wide, pairwise *F*_*ST*_ was calculated between all breed combinations to summarise overall differentiation. To locate differentiation along the genome, *F*_*ST*_ was additionally calculated in sliding windows (50 kb window, 25 kb step) between each Dutch tra- ditional breed and the Holstein Friesian. Windows containing fewer than 10 segregating sites were discarded, and windows in the top 0.1 % of the remaining distribution were retained as candidate regi- ons. Overlapping or adjacent candidate windows were merged into contiguous regions before counting.

Population structure was further investigated through a principal component analysis (PCA) using PLINK (v2.0.0a.6.9) (Chang et al., 2015), performed separately for SNPs, indels, and SVs. SNPs and indels were restricted to biallelic sites with half-calls set to missing, and pruned for linkage disequilibrium using a 50-variant window, a 5-variant step and an *r*^2^ threshold of 0.2. SVs were restricted to biallelic sites but were neither frequency-filtered nor pruned, as the marker set was too small for window-based pruning. All 53 principal components were extracted, so that reported variances are expressed relative to total variance. Ancestry proportions were estimated with ADMIXTURE (v1.4.0) (Alexander et al., 2009) for *K* = 2 to 10, run separately on the LD-pruned SNP and indel sets and on the unpruned biallelic SV set, selecting *K* by five-fold cross-validation error.

All visualisations, including *F*_*ST*_ heatmaps, windowed *F*_*ST*_ plots, and PCA plots, were generated in R (v4.5.2) (R Core Team, 2018) using the tidyverse ecosystem (v2.0.0) (Wickham et al., 2019), including the packages dplyr (v1.1.4) (Wickham et al., 2023b), ggplot2 (v4.0.1) (Wickham et al., 2023a), purrr (v1.2.0) (Henry and Wickham, 2023), readr (v2.1.6) (Wickham et al., 2023c), stringr (v1.6.0) (Wickham, 2022), and tibble (v3.3.0) (Müller and Wickham, 2023).

To identify SVs associated with breed-defining characteristics, the genotyped SV dataset was filtered based on population frequency and genomic location. An SV was defined as breed-specific if it was present in all short-read samples of the target breed (7/7 for traditional Dutch breeds and 5/5 for Holstein Friesian) and absent from all but at most one individual per non-target breed, with this tolerance applied to accommodate potential genotyping or sequencing errors. To provide statistical support for breed-specificity, *V*_*ST*_ (Redon et al., 2006) was computed for each SV between the target breed and all remaining animals from genotype dosages (0, 1 or 2 copies of the alternative allele), as *V*_*ST*_ = (*V*_*T*_ *™ V*_*S*_)*/V*_*T*_, where *V*_*T*_ is the total variance in dosage and *V*_*S*_ the sample-size-weighted mean within-group varia- nce. For each breed, *V*_*ST*_ was computed for all 55 865 SVs to obtain a genome-wide reference distribution, against which the empirical rank of each breed-specific SV was determined. Significance was asses- sed by permuting breed labels across animals (10 000 permutations), and *P*-values were adjusted with the Benjamini–Hochberg procedure. These calculations were performed in R (v4.5.2) (R Core Team, 2018) with the tidyverse (Wickham et al., 2019). Following this freque- ncy filtering, variants were cross-referenced with the bovine reference genome annotation (ARS-UCD2.0) by intersecting SV start coordi- nates with the GFF3-derived genomic features. This allowed for the identification of variants located within genic regions; genes affected were cross-referenced with their documented biological roles in cattle and with breed-specific phenotypes.

### Structural variant analysis

Structural variants were extracted from the pangenome short-read genotyping output and defined as alleles differing from the reference by at least 50 bp in length. We categorised SVs by type (insertions and deletions) and further classified them into four size classes: small (50 bp to 500 bp), medium (500 bp to 5 kb), large (5 kb to 50 kb), and very large (*>* 50 kb). SV coordinates were extracted using BCF- tools query and intersected with the ARS-UCD2.0 gene annotation (NCBI GCF 002263795.3 ARS-UCD2.0) to determine their genomic context. Where an SV overlapped more than one gene model, all overlapping genes were retained.

To identify regions of high genomic variation, we calculated SV and gene densities across the genome using non-overlapping 1 Mb win- dows, 10 Mb windows, and 25 Mb windows. The relationship between these densities was analysed to identify regions of high SV density.

## Results

### Sequencing data quality

For the short-read data, whole-genome sequencing yielded on average 30.0 Gb per sample (min 23.9 Gb, max 39.9 Gb), corresponding to an average genome coverage of 12.0 *×* (min 9.6 *×*, max 16.0 *×*) for the 54 animals. For the long-read data, reads from the 14 samples totalled on average 73.4 Gb per sample (min 21.5 Gb, max 112.7 Gb), with an average read N50 of 21.5 kb (min 6.9 kb, max 41.9 kb). The sample with the lowest yield, shortest reads and lowest read quality (21.5 Gb; N50 6.9 kb) produced an assembly that did not pass quality control.

### Genome assemblies & pangenome

We generated 14 *de novo* genome assemblies (Table 2), of which twe- lve assemblies exhibited high contiguity and completeness, with N50 values exceeding 100 Mb and complete BUSCO scores *>* 95 %. Two assemblies were excluded from pangenome construction due to quality constraints: DF 1 showed high fragmentation (N50 = 8.9 Mb), and DR 2 exhibited substantially lower gene-space completeness (BUSCO complete and single-copy = 88.8 %) in combination with lower N50 compared to the other assemblies.

### Pangenome graph composition

The final pangenome graph was constructed from 13 assemblies representing seven traditional Dutch cattle breeds (12 assemblies), supplemented by a Holstein Friesian assembly (PRJNA668863) where the ARS-UCD2.0 Hereford reference genome was used as backbone. The resulting graph comprises 138 019 429 nodes and 196 969 648 edges, with a mean node length of 20.9 bp.

We identified non-reference nodes defined as nodes not traversed by any ARS-UCD2.0 reference walk. This yielded 36 393 107 nodes containing 193.4 Mb of sequence representing variation encoded in alternative alleles, insertions, and assembly-specific expansions. Non-reference sequence content varied substantially across assemblies, with unique contributions ranging from 3.3 Mb to 37.5 Mb per genome, while shared non-reference sequence ranged from 12.8 Mb to 16.2 Mb per genome (approximately 0.15 % to 1.4 % of assembly length). Summed across both categories, individual assemblies contributed between 18.6 Mb and 50.3 Mb of non-reference sequence (Table 2),a nearly three-fold range within a panel of otherwise comparable contiguity and completeness. The largest contributors were the Meuse-Rhine-Yssel 3 (50.3 Mb) and Groningen White Headed 2 (44.3 Mb) samples, which are also the two longest *de novo* assemblies, while the Holstein Friesian assembly contributed the least of any genome in the panel (18.6 Mb).

**Table 2.** Comparison of the *de novo* generated genome assemblies of Dutch cattle breeds and the publicly available Holstein Friesian assembly (PRJNA668863). BUSCO scores consist of the following: complete and single-copy (S), complete and duplicated (D), fragmented (F), and missing (M), all in percentages of the total. Repeat percentage shows the percentage of bases masked by RepeatMasker. Non-reference sequence indicates the base pairs contributed as non-reference nodes in the pangenome graph, split in unique (only present in this assembly) and in shared (present in multiple assemblies, excluding ARS-UCD2.0). DBE: Dutch Belted, DF: Dutch Friesian, DFR: Dutch Friesian Red, DR: Deep Red, GWH: Groningen White Headed, HF: Holstein Friesian, IR: Improved Red, MRY: Meuse-Rhine-Yssel. Red indicates assemblies excluded from pangenome construction due to poor assembly quality, it also shows which statistics of the assembly led to their exclusion.

| Assembly | Length (Gb) | N50 (Mb) | BUSCO (%) | Repeats (%) | Unique non-ref (Mb) | Shared non-ref (Mb) |
|---|---|---|---|---|---|---|
| DBE | 2.644 | 103.7 | S:96.5,D:0.9,F:1.3,M:1.2 | 43.15 | 6.4 | 15.9 |
| DF 1 | 2.688 | 8.9 | S:96.5,D:1.1,F:1.5,M:0.9 | 43.88 | — | — |
| DF 2 | 2.642 | 103.9 | S:96.9,D:0.9,F:1.3,M:0.9 | 43.27 | 3.3 | 15.5 |
| DFR | 2.683 | 101.2 | S:96.4,D:1.1,F:1.5,M:1.0 | 43.79 | 15.8 | 14.6 |
| DR 1 | 2.645 | 103.7 | S:96.6,D:0.9,F:1.4,M:1.1 | 43.07 | 4.8 | 14.8 |
| DR 2 | 2.527 | 87.1 | S:88.8,D:0.8,F:5.1,M:5.2 | 41.94 | — | — |
| GWH 1 | 2.685 | 101.2 | S:96.5,D:1.1,F:1.5,M:0.9 | 44.09 | 18.8 | 13.0 |
| GWH 2 | 2.791 | 105.3 | S:95.6,D:1.3,F:1.7,M:1.3 | 43.75 | 30.4 | 13.9 |
| GWH 3 | 2.652 | 104.3 | S:96.8,D:0.9,F:1.4,M:0.9 | 43.61 | 3.5 | 16.2 |
| IR | 2.657 | 101.4 | S:96.6,D:1.0,F:1.3,M:1.1 | 43.51 | 5.9 | 14.0 |
| MRY 1 | 2.674 | 101.2 | S:96.5,D:1.0,F:1.4,M:1.1 | 43.82 | 7.8 | 13.4 |
| MRY 2 | 2.656 | 101.3 | S:96.2,D:1.0,F:1.4,M:1.4 | 43.48 | 12.9 | 13.6 |
| MRY 3 | 2.696 | 104.5 | S:96.1,D:1.0,F:1.7,M:1.2 | 43.73 | 37.5 | 12.8 |
| MRY 4 | 2.647 | 104.2 | S:96.7,D:0.9,F:1.3,M:1.0 | 43.25 | 3.7 | 15.8 |
| HF | 2.700 | 101.0 | — | — | 4.2 | 14.4 |
| ARS-UCD2.0 | 2.770 | 103.3 | S:97.5,D:1.3,F:0.5,M:0.6 | — | — | — |

We characterised the biological composition of non-reference sequence, by querying nodes of at least 200 bp (156 148 nodes) against the ARS-UCD2.0 bovine RNA database and the Dfam repeat library. Fewer than a third (30.9 %) of the nodes returned a match to either database: 25.1 % of nodes overlapped a known repeat element and 21.9 % showed homology to an annotated bovine transcript, with 16.2 % matching both, consistent with transposable element insertions within gene bodies or repeat-associated gene duplications. Among repeat matches, LINEs predominated (22.7 % of all queried nodes), with LTR elements (1.6 %) and SINEs (0.8 %) contributing far less. Transcript matches were mainly mRNA (13.4 %) and lncRNA (7.2 %). The remaining 69.1 % of nodes *≥*200 bp matched neither database, representing candidate novel sequence not captured by current bovine annotations.

### Non-reference sequence is predominantly breed-private

Analysis of non-reference node sharing across breeds revealed that the majority of non-reference sequence in the pangenome is breedspecific, and that this asymmetry becomes more pronounced with increasing variant size (Table S1; Figure 1).Meuse-Rhine-Yssel and Groningen White Headed carried the largest amounts of breed-private sequence overall, substantially exceeding all other breeds; these are also the only breeds contributing multiple assemblies (four and three respectively). This excess was concentrated almost entirely in the largest size class: nodes *≥*1 kb contributed more than the entire breed-private total of other breeds.

**Figure 1.**
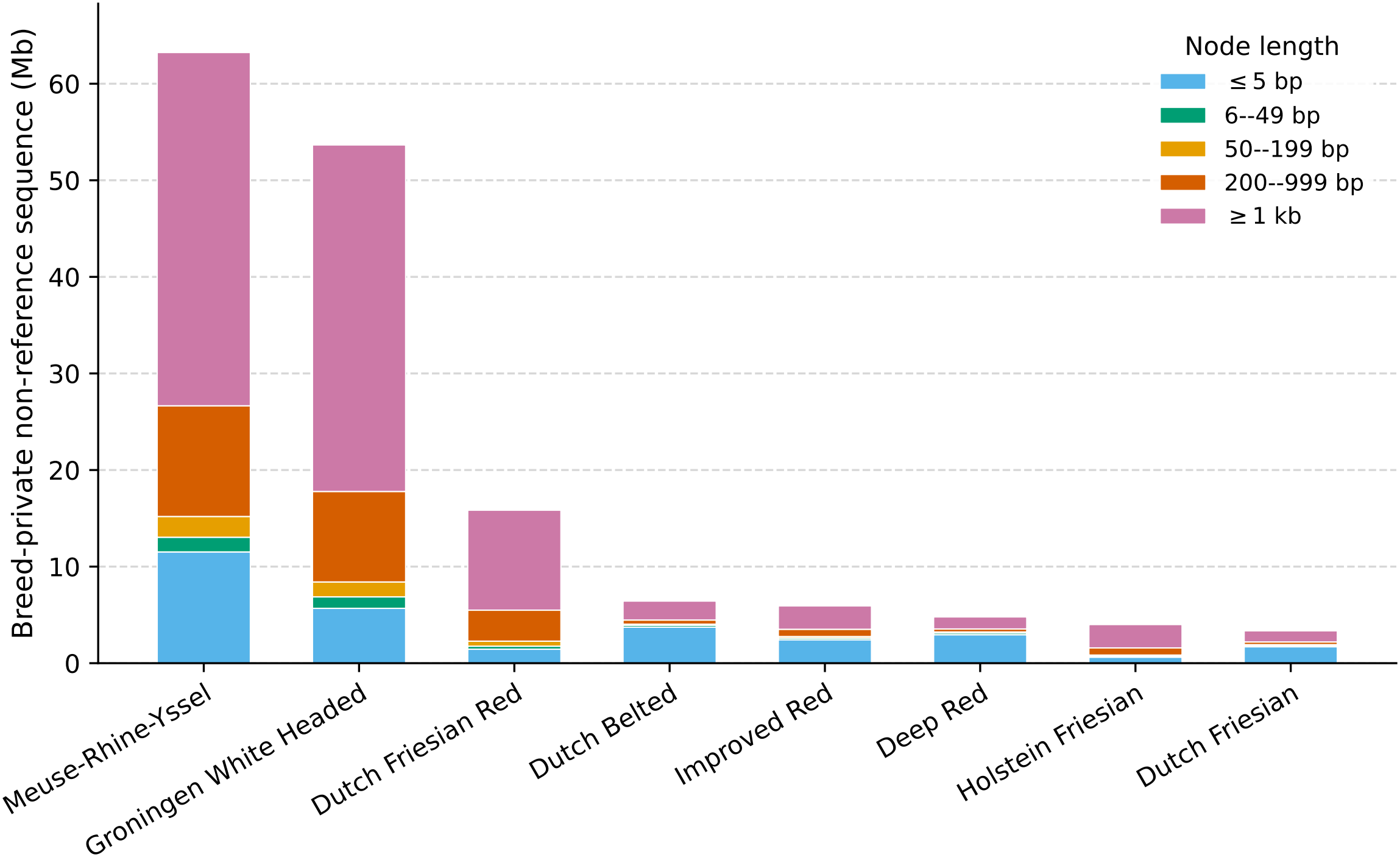
Non-reference sequence carried by each breed is dominated by long, breed-private nodes. Breed-private non-reference sequence in the Dutch cattle pangenome, stratified by node length. Bars represent the total sequence (Mb) in non-reference graph nodes exclusively traversed by assemblies of a single breed, partitioned into five length strata: SNP-level (*≤*5 bp), small indel (6 bp to 49 bp), short SV (50 bp to 199 bp), intermediate SV (200 bp to 999 bp) and long SV (*≥*1 kb).

All pangenome non-reference nodes were classified into one of the three categories, breed-private (traversed by assemblies of a single breed only), multi-breed shared (traversed by assemblies of more than one but not all breeds), or pan-breed (traversed by at least one assembly of every breed), and further stratified into five length bins (Table S2).

Breed-private sequence dominated the non-reference component of the graph overall; this dominance was strongly size-dependent as the fraction of non-reference sequences classified as breed-private was much higher for extremely long structural variants (*≥*1 kb) than for small-indels (*≤*5 bp). In terms of node counts, breed-private nodes outnumbered multi-breed shared and pan-breed nodes combined at every length stratum except the indel classes 6 bp to 49 bp and 50 bp to 199 bp, where multi-breed shared nodes were more numerous, despite contributing less total sequence.

Because breeds represented by several assemblies can accumulate breed-private sequence purely through assembly number, all totals were recomputed after downsampling to one assembly per breed. This reduced the non-reference sequence to 83.79 Mb, of which 54.59 Mb was breed-private, 26.78 Mb multi-breed shared, and 2.42 Mb pan-breed. The breed-private fraction fell from 81.3 % to 65.2 %, but the size dependence was unchanged; breed-private sequence still accounted for 98.3 % of nodes *≥*1 kb (Table S3).

### Pangenome quality validation against known variants

Having characterized the general composition of the pangenome, we next tested whether the graph could recover known functional variants through two benchmarks: the recovery of a known large copy number variant (CNV) in the Dutch Belted and the genotyping accuracy of breed-specific SNPs in the Groningen White Headed.

#### SV validation

We investigated the CNV on BTA3 connected the belted phenotype in Dutch Belted cattle, earlier analysed by Awasthi Mishra et al. (2017), Rothammer et al. (2018), and Hogers et al. (2026). The belted phenotype is driven by a complex CNV on BTA3, with non-belted animals carrying a single copy and belted individuals commonly four tandem copies per locus (Awasthi Mishra et al., 2017; Rothammer et al., 2018). The graph encoded this locus as a single large insertion (Figure S1), capturing its presence and approximate scale but collapsing the tandem copies onto one alternate path rather than as parallel, countable paths. Because k-mer genotypers such as PanGenie resolve genotypes by comparing coverage across alternative paths through a bubble, no local structure exists against which copy number can be inferred, and short-read genotyping accordingly failed to resolve copy number in Dutch Belted samples. Pangenomes can therefore represent large, complex SVs that a linear reference cannot capture at all, but their utility for genotyping copy number depends on how the graph is decomposed during construction.

#### SNP validation

We next analysed nine SNPs on BTA20 known to be strongly associated with the Groningen White Headed breed (Gonzalez-Prendes et al., 2022). First, we assessed their presence within the *de novo* assemblies used to build the graph, finding that all SNPs were present in at least one of the three Groningen White Headed assemblies (Table S4). Within the short-read cohort, six of the nine markers reached fixation in the Groningen White Headed samples. However, a notable discrepancy was observed at position 20 044 910 bp. While present in two Groningen White Headed assemblies, the variant was absent in all Groningen White Headed short-read samples and was instead detected in a single Dutch Friesian individual. Further analysis showed that at this position no SNP was found, but instead it featured a multiple nucleotide polymorphism of two bp. Where these markers appeared in other breeds they were invariably heterozygous, whereas Groningen White Headed carriers were usually homozygous. To place these genotypes in the context of a non-pangenome work-flow, the same 54 animals were independently genotyped against the linear ARS-UCD2.0 reference at the nine marker positions. Overall agreement was high, with 471 of 486 comparisons concordant (96.9 %) and six of the nine markers in complete agreement. The 15 discordant calls were, however, strongly directional: in 14 cases the linear call carried more copies of the alternate allele than the pangenome call, and 13 involved Groningen White Headed animals. At 20 044 910 bp the linear reference recovered the alternate allele in all seven Groningen White Headed individuals, six of them homozygous, whereas pangenome genotyping called each as homozygous reference. The two remaining discordant markers showed the same directional pattern at lower magnitude (20 044 595 bp, four Groningen White Headed animals under-called, 20 542 032 bp, two called heterozygous where linear calling gave the homozygous alternate).

### Genetic diversity in Dutch cattle breeds

To characterise genetic diversity among the traditional Dutch cattle breeds, we analysed the 54 short-read sequenced animals (Table 1).Initial variant calling across the pangenome yielded a total of 46 668 791 raw variants, comprising 17 766 196 SNPs, 24 467 862 insertions and deletions, 2 598 950 multi-nucleotide polymorphisms, and 1 835 783 complex alleles. Monomorphic variants were removed (30 991 064; 66.4 % of the raw set), of which 30 848 737 were homozygous reference and 142 327 homozygous alternative. Variants on the sex chromosomes and mitochondrial genome were also excluded (2 926 147 X-linked, 143 725 Y-linked, and 301 mitochondrial). In total, we retained 15 235 145 polymorphic autosomal variants: 10 750 422 SNPs, 3 676 817 insertions and deletions, 597 407 multinucleotide polymorphisms, and 210 499 complex alleles. Filtering disproportionately affected indels, of which 85.0 % were removed compared with 39.5 % of SNPs. Genotype missingness in the retained set was negligible (84 246 missing calls out of 822 697 830; 0.010 %).

Within-breed diversity was quantified on the biallelic autosomal SNP set (Table 3).Nucleotide diversity was lowest in the Groningen White Headed (*π* = 0.001 18) and highest in the Improved Red (0.001 40), with the remaining breeds spanning a narrow range in between; Holstein Friesian fell mid-range (0.001 30). Individual heterozygosity followed a similar ordering, from 958 heterozygous sites per Mb of autosome in the Dutch Friesian to 1198 in the Improved Red. Runs of homozygosity gave the reverse: mean *F*_*ROH*_ was highest in the Dutch Friesian (0.355) and the Groningen White Headed (0.352), and lowest in the Improved Red (0.225) and Deep Red (0.255). Restricted to segments *≥*1 Mb, which reflect more recent inbreeding, the Groningen White Headed ranked highest (0.084) and the Improved Red lowest (0.010).

**Table 3.**
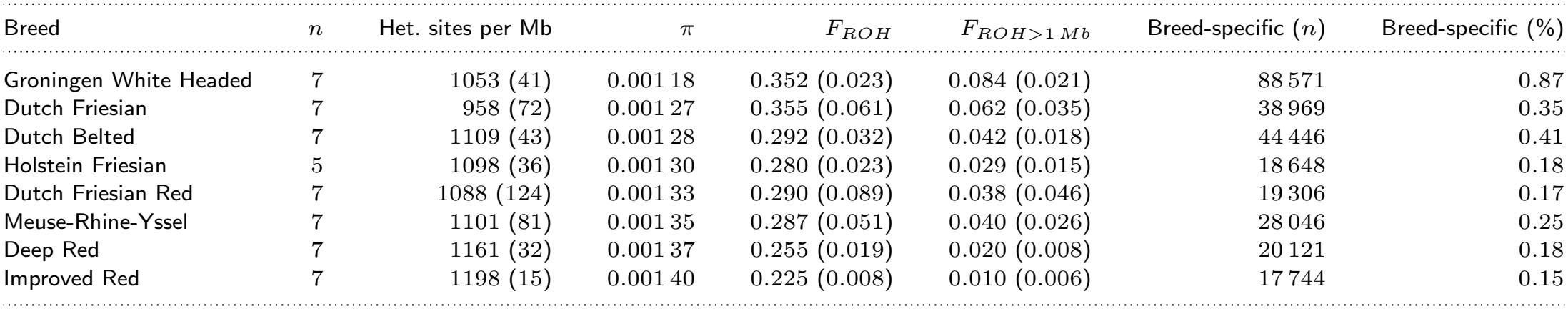
Within-breed genetic diversity, estimated from biallelic autosomal SNPs. Heterozygous sites per Mb are per individual, scaled to the 2 489 385 779 bp ARS-UCD2.0 autosomal genome. *π* is the mean of 100 kb windows. *F*_*ROH*_ is the genome fraction in runs of homozygosity, over all segments called; because no minimum segment length was imposed these values include short tracts reflecting background linkage disequilibrium and are informative for ranking breeds rather than as absolute autozygosity. *F*_*ROH>*1 *Mb*_ includes only segments *≥*1 Mb, reflecting recent autozygosity. Breed-specific variants are those present in at least two individuals of the focal breed and absent from all individuals of every other breed, with the percentage expressed relative to the variants segregating within that breed. Standard deviations across individuals are given in parentheses. Breeds are ordered by *π*.

| Breed | *n* | Het. sites per Mb | π | *F*<sub>*ROH*</sub> | *F*<sub>*ROH*>1 *Mb*</sub> | Breed-specific (*n*) | Breed-specific (%) |
|---|---|---|---|---|---|---|---|
| Groningen White Headed | 7 | 1053 (41) | 0.001 18 | 0.352 (0.023) | 0.084 (0.021) | 88 571 | 0.87 |
| Dutch Friesian | 7 | 958 (72) | 0.001 27 | 0.355 (0.061) | 0.062 (0.035) | 38 969 | 0.35 |
| Dutch Belted | 7 | 1109 (43) | 0.001 28 | 0.292 (0.032) | 0.042 (0.018) | 44 446 | 0.41 |
| Holstein Friesian | 5 | 1098 (36) | 0.001 30 | 0.280 (0.023) | 0.029 (0.015) | 18 648 | 0.18 |
| Dutch Friesian Red | 7 | 1088 (124) | 0.001 33 | 0.290 (0.089) | 0.038 (0.046) | 19 306 | 0.17 |
| Meuse-Rhine-Yssel | 7 | 1101 (81) | 0.001 35 | 0.287 (0.051) | 0.040 (0.026) | 28 046 | 0.25 |
| Deep Red | 7 | 1161 (32) | 0.001 37 | 0.255 (0.019) | 0.020 (0.008) | 20 121 | 0.18 |
| Improved Red | 7 | 1198 (15) | 0.001 40 | 0.225 (0.008) | 0.010 (0.006) | 17 744 | 0.15 |

We next examined breed-specific variants, defined as variants found in at least two individuals of a single breed and absent from all others (Table 3).The Groningen White Headed carried by far the largest share (88 571 variants; 0.87 % of its segregating sites), followed by the Dutch Belted (0.41 %) and the Dutch Friesian (0.35 %); the lowest group consisted of Meuse-Rhine-Yssel (0.25 %), Deep Red, Dutch Friesian Red, and Improved Red (0.15 % to 0.18 %). In comparison the commercial Holstein Friesian was among the lowest (0.18 %).

Genome-wide pairwise *F*_*ST*_ was used to summarise the overall level of differentiation between breeds (Figure S2).The Groningen White Headed emerged as the most genetically distinct population, showing the highest differentiation from the Dutch Belted and Dutch Friesian. Conversely, the lowest differentiation was observed within the red-breed group, between Deep Red and both Meuse-Rhine-Yssel and Improved Red (0.017 in each case), followed by the two Friesian breeds (0.025).

Genome-wide averages describe the mean level of divergence but not its distribution along the genome. To identify localised regions of elevated differentiation, which may reflect breed-specific selection or introgression, we additionally calculated *F*_*ST*_ in 50 kb windows (Figure S3) between each of the seven Dutch traditional breeds and Holstein Friesian. As expected, windowed *F*_*ST*_ values were higher and considerably more variable than the genome-wide estimates. The genome-wide value is a ratio of averages across all sites, whereas the mean of per-window values is an average of ratios and is upwardly biased when few segregating sites contribute to each window (Bhatia et al., 2013); windows were therefore ranked within each comparison rather than compared against the genome-wide values on an absolute scale. An elevated cluster of windows on BTA20 appeared in six of the seven breed comparisons, ranking as the top or second-ranked chromosome in four of them (Deep Red, 20 windows; Dutch Friesian, 15; Groningen White Headed and Meuse-Rhine-Yssel, 12 each; Improved Red, 10; Dutch Friesian Red, 4), indicating a shared candidate region of differentiation between Holstein Friesian and multiple, independently differentiated Dutch traditional breeds. Cross-examining the windowed *F*_*ST*_ coordinates identified 12 distinct BTA20 windows shared by at least two breeds (Appendix, Table S5), the most conserved lying at 42.32 Mb to 42.37 Mb and surfacing in four populations. The Dutch Belted was the sole exception, with no BTA20 window reaching the top 0.1 % threshold; its differentiation was instead concentrated overwhelmingly on BTA3 (Figure 2), which contributed 27 of the 91 windows above the threshold, more than three times the next-ranked chromosome. These windows formed three clusters, at approximately 104.6 Mb to 105.8 Mb, 111.4 Mb to 114.0 Mb and 116.7 Mb to 118.1 Mb, the strongest signal genome-wide falling at 117.85 Mb to 117.90 Mb (*F*_*ST*_ = 0.823, 421 segregating sites). The distal cluster contains the copy number variant underlying the belted phenotype at 118 028 480 bp to 118 034 892 bp (Hogers et al., 2026), which is bracketed by two windows in the top 0.1 % (*F*_*ST*_ = 0.658 ending 3.5 kb proximal to the variant and 0.728 beginning 15.6 kb distal to it). The two windows spanning the variant itself fell just short of the threshold (*F*_*ST*_ = 0.624 and 0.586) and contained 54 and 80 segregating sites, against 134 and 148 in the flanking windows and 300 to 425 in the most differentiated windows of the cluster.

**Figure 2.**
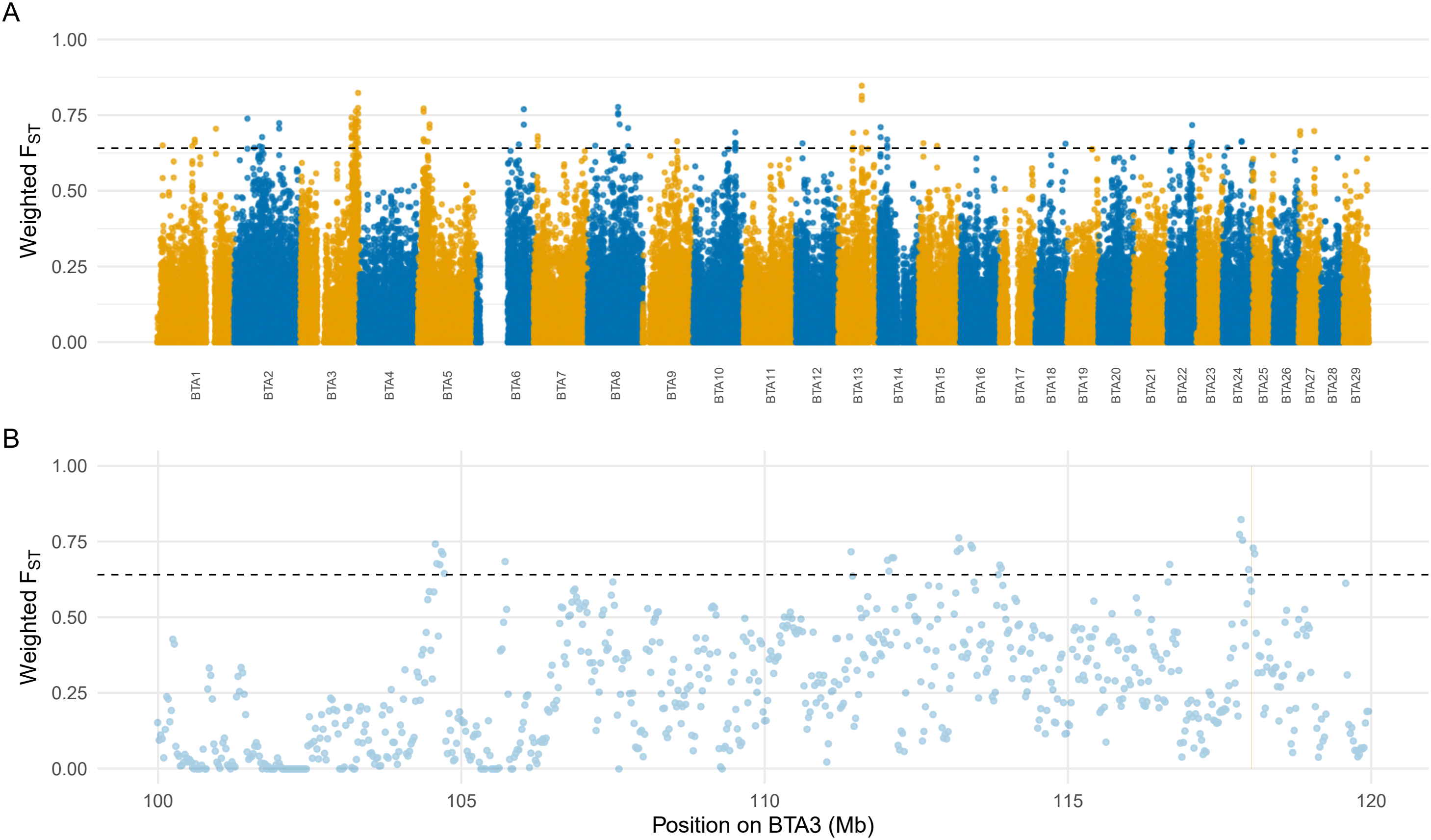
Genomic differentiation between the Dutch Belted and Holstein Friesian is concentrated on BTA3. Windowed pairwise *F*_*ST*_ (50 kb windows, 25 kb step, Weir and Cockerham estimator) between the Dutch Belted and Holstein Friesian. (A) Genome-wide, with points coloured by chromosome (BTA1–BTA29); the horizontal dashed line marks the top 0.1 % empirical threshold. (B) The BTA3 region between 100 Mb to 120 Mb, containing the three clusters of elevated differentiation; the shaded band marks the copy number variant underlying the belted phenotype at 118 028 480 bp to 118 034 892 bp (Hogers et al., 2026).

Principal component analysis (PCA) further resolved the genetic relationships between the breeds (Figure 3). For SNPs, PC1 primarily separated the Groningen White Headed from the remaining breeds, whereas PC2 distinguished the Dutch Friesian, Dutch Friesian Red, and Dutch Belted populations. PC3 further isolated the Dutch Belted, while PC4 primarily separated the Holstein Friesian from the traditional Dutch breeds. Indels and SVs broadly recapitulated this structure, however, the separation among breeds for SVs was generally less distinct than for SNPs and indels, and a small number of Dutch Friesian individuals appeared as outliers along PC2 and PC4. Dutch Belted, Holstein Friesian, and Groningen White Headed individuals consistently formed separate clusters across all three variant types, differentiating them from the other breeds. Scree plots for all three variant classes are provided in the Appendix (Figure S4), with variance explained declining steeply through PC4 before flattening.

**Figure 3.**
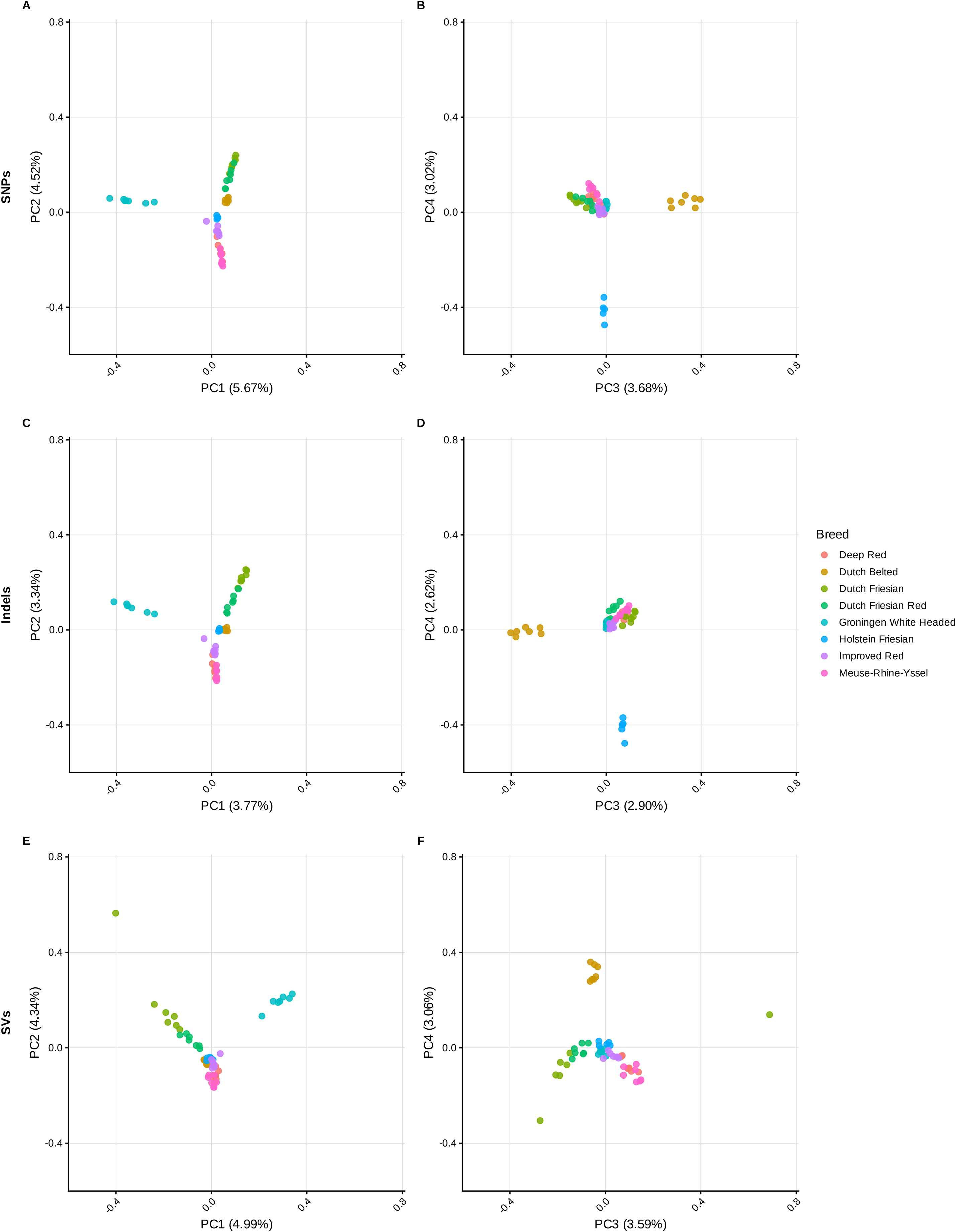
SNPs, indels and structural variants resolve the same four population clusters. Principal component analysis (PCA) of eight Dutch cattle populations based on SNPs (A, B), indels (C, D), and structural variants (E, F). Panels show PC1 versus PC2 (left column) and PC3 versus PC4 (right column) for each variant class.

Model-based clustering with ADMIXTURE was applied to the same variant sets. Cross-validation error was minimised at *K* = 2 for all three variant classes (0.631 for SNPs, 0.559 for indels, 0.536 for SVs), separating the Groningen White Headed from all other populations. Higher values of *K* resolved the red breeds, the Friesian breeds and the Dutch Belted as separate components, consistent with the structure recovered by PCA, and the three variant classes gave concordant results (Figure S5).

### Structural variants landscape in Dutch cattle

One of the primary advantages of utilising pangenomes is the inclusion of structural variants, enabling the genotyping of large-scale genetic variation using short-read data. A total of 55 865 structural variants (*≥* 50 bp) were identified in our 54 sample panel, comprising 32 764 insertions and 23 101 deletions. The insertions add 28.28 Mb of sequence absent from ARS-UCD2.0, while the deletions remove 19.16 Mb of reference sequence. Most SVs were on the smaller side, 39 180 variants fell between 50 bp and 500 bp (70.1 %), 14 944 between 500 bp and 5 kb, 1713 between 5 kb and 50 kb, and only 28 exceeded 50 kb.

Consistent with previous observations of SV landscapes, smaller structural variants were more abundant (Figure 4);this includes an increased concentration of variants between 100 bp to 400 bp, corresponding to the expected size of Short Interspersed Nuclear Elements (SINEs), and similarly, an uptick was observed near 6 kb, corresponding to the expected size of Long Interspersed Nuclear Elements (LINEs). Interestingly, while insertions and deletions appear in roughly equal proportions for smaller SVs, the count of deletions drops sharply starting at approximately 500 bp, whereas insertions decrease at a much slower rate. From 5 kb onwards, insertions are markedly more frequent than deletions; at the 10 kb threshold, insertions represent nearly all identified SVs before the distribution levels out for very large variants (*>* 50 kb).

**Figure 4.**
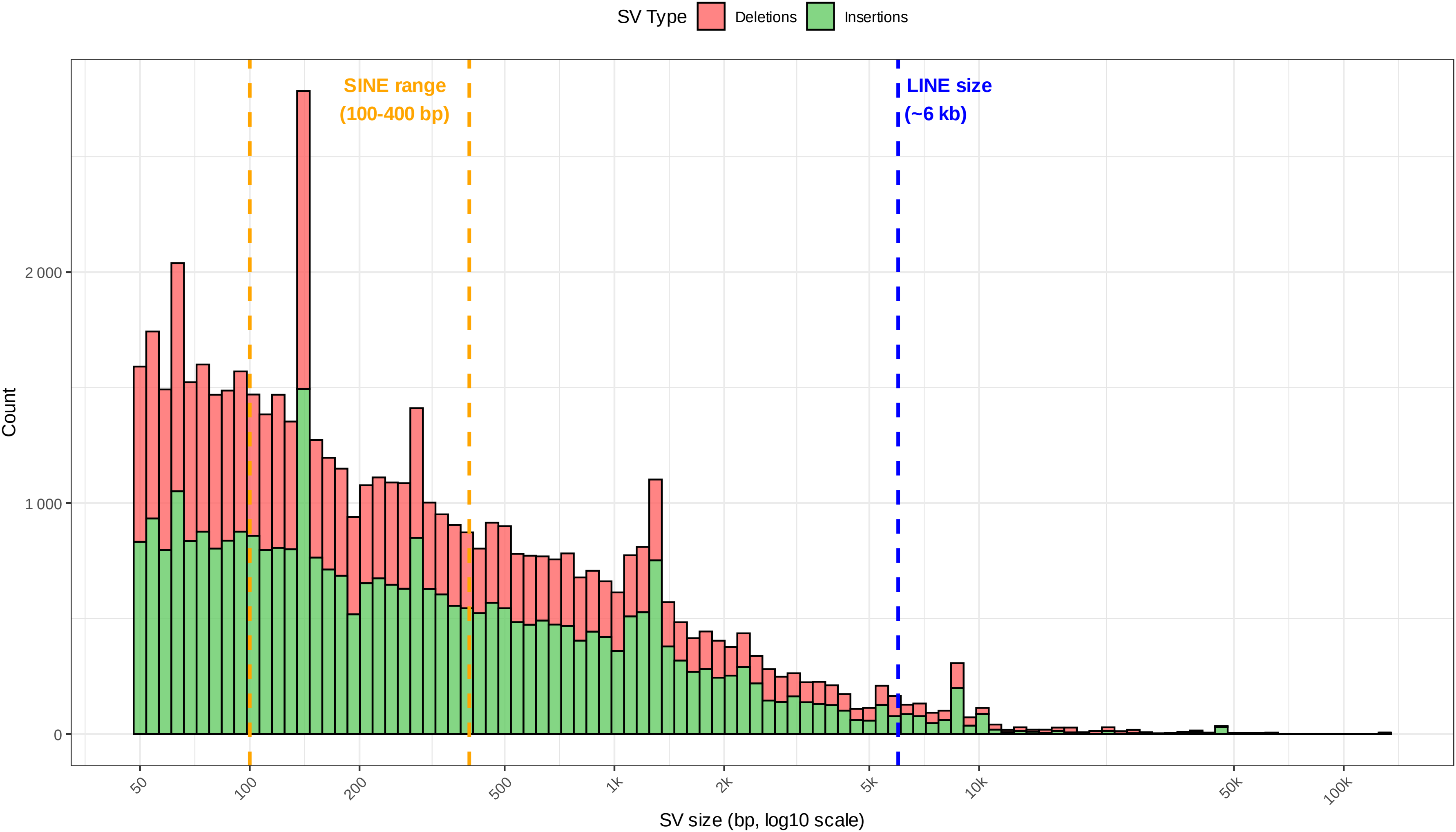
Structural variant sizes carry a mobile-element signature at SINE and LINE lengths. Size distribution of structural variants (50 bp to 1 Mb). The histogram highlights the prevalence of Short Interspersed Nuclear Element (SINEs, 100 bp to 400 bp) and Long Interspersed Nuclear Elements (LINEs, 6 kb).

The genomic density of larger variants is often hypothesised to be inversely related to gene density, as large-scale mutations are more likely to disrupt coding sequences, affect splice sites, or interrupt transcription, thereby theoretically facing stronger purifying selection. We correlated SV density against both gene count and gene nucleotide coverage in 1 Mb, 10 Mb, and 25 Mb bins. All correlations were negative but negligible in magnitude, explaining at most 12 % and typically under 2 % of the variance in SV density (*R*^2^ = 0.0002 to 0.123). Structural variant density in this dataset is therefore not appreciably predicted by gene content.

To investigate the potential genetic drivers of breed-defining traits, SVs were filtered to identify those consistently present within a specific population. An SV was required to be found in all seven individuals of a target breed, while being present in no more than one individual per breed for the other breeds. Following these criteria, 10 fixed breed-specific SVs were identified within genic regions (Table 4).All ten ranked within the top 0.15 % of the genome-wide *V*_*ST*_ distribution of their breed (nine within the top 0.1 %; *V*_*ST*_ = 0.52 to 0.90), and all were significant in the permutation test after correction for multiple testing (*q <* 1 *×* 10^*−*4^, the minimum attainable with 10 000 permutations). Genome-wide, the top 0.1 % *V*_*ST*_ threshold was highest for the Groningen White Headed (0.685) and lowest for the Improved Red (0.295), consistent with the SNP-based *F*_*ST*_ results.

**Table 4.** Summary of breed-specific structural variants (SVs) by gene, chromosome, position, length, type, and breed breakdown. *V*_*ST*_ is computed between the target breed and all other animals from genotype dosages; Top (%) is the percentage of all 55 865 SVs with an equal or higher *V*_*ST*_ in the same comparison. All SVs had a Benjamini–Hochberg-adjusted permutation *q <* 1 *×* 10^*−*4^ (10 000 permutations). Breed abbreviations: DBE: Dutch Belted, DF: Dutch Friesian, DFR: Dutch Friesian Red, DR: Deep Red, GWH: Groningen White Headed, IR: Improved Red, MRY: Meuse-Rhine-Yssel, HF: Holstein Friesian.

| Chromosome | Start position (bp) | Gene | SV length (bp) | SV type | Breed Breakdown | *V*<sub>*ST*</sub> | Top (%) |
|---|---|---|---|---|---|---|---|
| BTA6 | 70 203 395 | *KIT* | 174 bp | Deletion | 7/7 DBE | 0.898 | 0.002 |
| BTA6 | 71 507 426 | *CRACD* | 50 bp | Insertion | 7/7 MRY; 1/7 DR; 1/7 DFR; 1/5 HF; 1/7 IR | 0.518 | 0.016 |
| BTA9 | 73 527 128 | *AHI1* | 96 bp | Insertion | 7/7 GWH; 1/7 IR | 0.856 | 0.020 |
| BTA11 | 102 909 039 | *AK8* | 518 bp | Insertion | 7/7 GWH | 0.898 | 0.013 |
| BTA11 | 102 909 097 | *AK8* | 184 bp | Insertion | 7/7 GWH | 0.898 | 0.013 |
| BTA16 | 76 527 021 | *DENND1B* | 66 bp | Insertion | 7/7 GWH | 0.903 | 0.009 |
| BTA16 | 76 527 928 | *DENND1B* | 66 bp | Deletion | 7/7 GWH | 0.903 | 0.009 |
| BTA16 | 76 528 014 | *DENND1B* | 654 bp | Deletion | 7/7 GWH; 1/7 DBE; 1/5 HF; 1/7 IR | 0.659 | 0.143 |
| BTA20 | 37 118 424 | *CPLANE1* | 1376 bp | Insertion | 7/7 GWH; 1/7 DFR; 1/7 MRY | 0.727 | 0.084 |
| BTA22 | 41 200 908 | *FHIT* | 98 bp | Deletion | 5/5 HF; 1/7 GWH; 1/7 DR; 1/7 IR; 1/7 MRY | 0.617 | 0.021 |

The Dutch Belted breed provided a clear link between genomic variation and phenotype through a 174 bp deletion on BTA6 within the *KIT* gene. This variant was found in all Dutch Belted individuals and was absent in all others. The *KIT* locus is a well-documented master regulator of melanocyte development, mutations in this gene are associated with various white-spotting and belting patterns across mammalian species (Giuffra et al., 1999; Andersson, 2020). The deletion had the highest *V*_*ST*_ (0.898) of all 55 865 SVs in the Dutch Belted comparison.

The Groningen White Headed exhibited the highest number of breed-specific SVs, further solidifying its genetic distinctiveness observed in the *F*_*ST*_ and PCA analyses. The largest was a 1376 bp insertion in *CPLANE1* (Ciliogenesis and Planar Polarity Effector Complex Subunit 1) on BTA20, fixed in the Groningen White Headed and found in only one Dutch Friesian Red and one Meuse-Rhine-Yssel individual. Furthermore, the Groningen White Headed possessed exclusive insertions in *AK8* (BTA11) and *DENND1B* (BTA16), as well as a specific insertion in *AHI1* (BTA9), which, like *CPLANE1*, is involved in ciliary assembly and signal transduction. Notably, the fixed *DENND1B* variants comprise a 66 bp insertion at position 76 527 021 and a 66 bp deletion of near-identical sequence 907 bp downstream at position 76 527 928.

Other breeds also displayed fixed variation within functional genes, though often with slightly more shared variation across the cohort. The MRY reached fixation for a 50 bp insertion in *CRACD* on BTA6, a gene associated with cell adhesion and cytoskeletal organisation. Interestingly, the Holstein Friesian outgroup exhibited fixation for a 98 bp deletion in *FHIT* (Fragile Histidine Triad), which appeared only sporadically in other breeds. These two SVs, each carried by four animals of other breeds, had the lowest *V*_*ST*_ of the set (0.52 and 0.62) but still ranked within the top 0.03 % of their breed comparisons.

## Discussion

Pangenomes are increasingly used in cattle, but the work so far has concentrated either on the technical merits of different graph construction approaches (Leonard et al., 2022; Smith et al., 2023; Leonard et al., 2023, 2024; Liu, 2025) or on single globally dominant breeds such as the Holstein Friesian (Yang et al., 2026). Traditional European populations, which are the least likely to be chosen as reference individuals and the most likely to disappear, have been largely absent. The Dutch cattle pangenome presented here addresses that gap, and the results support three main conclusions: that Dutch traditional breeds carry a large body of non-reference sequence, most of it breed-private, that a linear reference cannot represent; that a single graph makes SNPs, indels and structural variants accessible to short-read genotyping in one pass, with the largest gain in the structural component; and that this combination opens a practical route to trait discovery and conservation genetics in populations too small to justify a dedicated reference genome each.

### Dutch traditional breeds carry a large, largely breed-private non-reference genome component

The pangenome graph contains 193.4 Mb of sequence not traversed by the ARS-UCD2.0 reference walk, exceeding the 116 Mb reported for a global cattle pangenome (Talenti et al., 2022) but well below the 552 Mb reported across 11 bovine species by Leonard et al. (2023), as expected for a panel of solely taurine breeds sampled within one country. However, breed selection is not the only factor in determining the size of the pangenome; graph builder and filtering criteria also play a major factor. Differences of almost a factor of five have been reported between Cactus and Minigraph (Leonard et al., 2023), Talenti et al. (2022) retained 116.1 Mb of an initial 257.2 Mb after removing misassembled regions and short variants, and in pigs 136.6 Mb of unaligned sequence was reduced to 72.5 Mb once sequences below 300 bp were discarded (Tian et al., 2020). Our assemblies were built from R9.4.1 ONT data and polished with short reads, the base-level error of that chemistry likely introduced spurious short variants and probably inflates our total, and also contributed to the 69.1 % of nodes *≥*200 bp that matched neither the bovine transcript set nor the Dfam repeat library. That fraction is not solely an error signal, however, intergenic non-repetitive sequence has no representation in a transcript database, and graph nodes are fragments rather than complete elements, so interior segments of a repeat frequently fail to align.

The striking feature of this non-reference component is how little of it is shared. Breed-private sequence accounts for 157.2 Mb (81.3 %) of the total, and the private fraction rises steeply with size, from 67.0 % at the SNP level to 99.3 % for nodes *≥*1 kb. Part of this is an artefact of panel design, since six of the eight breeds are represented by a single assembly and a rare large variant is unlikely to be sampled twice. Downsampling to one assembly per breed lowers the overall private fraction substantially, to 65.2 %, but leaves the size dependence intact: nodes *≥*1 kb remain 98.3 % breed-private. It is likely that short variants persist across lineages through shared ancestry or recurrent mutation, whereas large insertions arise in recent, lineage-specific events with little opportunity to spread between breeds. The Meuse-Rhine-Yssel and the Groningen White Headed carried the most breed-private sequence (63.2 Mb and 53.7 Mb), but they are also the only breeds with multiple assemblies, and after downsampling the Dutch Friesian Red ranks highest (17.08 Mb); assembly number, not biology, largely drove the raw ordering.

Individual assemblies contributed between 18.6 Mb and 50.3 Mb of non-reference sequence, a nearly three-fold range within a panel of otherwise comparable quality. The largest contributors, Gronin-gen White Headed 2 (44.3 Mb), and Meuse-Rhine-Yssel 3 (50.3 Mb), are also the largest and most contiguous assemblies, so some of their excess is likely residual assembly artefact rather than biological sequence. At the other extreme, the Holstein Friesian assembly contributed the least of any genome in the panel. Part of this difference likely reflects assembly strategy, since the Holstein Friesian genome was generated independently and is expected to carry fewer residual base-level errors than our ONT R9.4.1 assemblies. Even so, that a single globally dominant commercial breed adds less novel sequence than any individual traditional Dutch assembly is a clear illustration of what is at stake in the loss of traditional populations.

### Graph-based genotyping reproduces known population structure and highlights the vulnerability of the Groningen White Headed

Genotyping the short-read cohort against the graph resolved four coherent groups: the Groningen White Headed, a red-breed cluster of Meuse-Rhine-Yssel, Deep Red, and Improved Red, a Friesian cluster of Dutch Friesian and Dutch Friesian Red, and the Dutch Belted alone. This structure is consistent across PCA and ADMIX-TURE, and largely agrees with earlier SNP-array characterisations of these populations (Hulsegge et al., 2017; van Breukelen et al., 2019; Marjanovic et al., 2021; Schmidtmann et al., 2021),indicating that genotypes called against the graph capture established breed relationships. The exception is the Dutch Belted, which formed its own cluster here but grouped with the Friesian breeds in earlier studies, with the Dutch Friesian in a relatedness analysis (Marjanovic et al.,2021) and with the Dutch Friesian Red in a population tree (Schmidtmann et al., 2021); along PC2 it also fell with the Friesian breeds in our data. Holstein Friesian was the only population without a cluster of its own, distributing across the red and Friesian components (0.53 and 0.27) and carrying one of the lowest proportions of breed-specific variants (0.18 %). This is what its history in the Netherlands predicts: Holsteinisation, which began around 1970, has left the Holstein Friesian at about 99 % of the registered female breeding cattle (Theunissen, 2012; Centrum voor Genetische Bronnen, 2026), and the grading-up of native cows with Holstein Friesian bulls has seeded Holstein Friesian haplotypes throughout the other breeds (van Breukelen et al., 2019; Hulsegge et al., 2022). At the opposite end, the low breed-specific variation of the Improved Red and Deep Red reflects recency rather than admixture: both were derived from the Meuse-Rhine-Yssel with herdbooks established only in 1988 and 2001, and the pairwise *F*_*ST*_ of 0.017 between the Deep Red and both its progenitor and the Improved Red leaves little room for accumulated drift (van Breukelen et al., 2019). The two Friesian breeds are similarly close (0.025), consistent with the origin of the Dutch Friesian Red within the Frisian population and the frequent exchange of bulls between the two breeds (Strikwerda, 2014; van Breukelen et al., 2019).

The Groningen White Headed is the most differentiated population by every measure applied here: the highest breed-specific variant proportion (0.87 %), the highest pairwise *F*_*ST*_ against every other breed, separation along PC1, and its own ADMIXTURE component at every *K*. However, it also carries the lowest nucleotide diversity in the panel (*π* = 0.001 18), the second-lowest individual heterozygosity, among the highest *F*_*ROH*_ (0.352), and the highest *F*_*ROH*_ in segments *≥*1 Mb (0.084; Table 3). Divergence and diversity are pulling in opposite directions, which is exactly what is expected of a small, historically isolated population under strong drift. At the start of the twentieth century the breed made up 90 % of all cattle in the province of Groningen, and from the mid-1970s most farmers crossed it with Holstein Friesian to raise milk production (Hiemstra et al., 2010). The Groningen White Headed holds a disproportionate share of the unique variation in this panel while having relatively little variation left within itself, so the diversity it represents is both especially valuable and especially fragile. The Dutch Friesian shows a milder version of the same profile (*F*_*ROH*_ = 0.355, the lowest individual heterozygosity in the panel), whereas the recently derived Improved Red and Deep Red retain the highest within-breed diversity despite their small census sizes, having inherited it from the much larger Meuse-Rhine-Yssel.

### The pangenome recovers structural variation that reference-based short-read approaches miss, including a fixed deletion in *KIT*

Graph-based genotyping of the short-read cohort yielded 55 865 structural variants, comprising 32 764 insertions adding 28.28 Mb of sequence absent from ARS-UCD2.0 and 23 101 deletions removing 19.16 Mb of reference sequence. The size spectrum carries a clear mobile-element signature, with a concentration between 100 bp to 400 bp matching SINEs and a secondary rise near 6 kb matching LINEs (Figure 4), consistent with the repeat composition of the long non-reference nodes, of which 22.7 % matched LINEs and 16.2 % matched both a repeat and an annotated transcript. Insertions and deletions occur in roughly equal numbers below 500 bp, but deletions fall away sharply above that threshold while insertions decline slowly, so that beyond 5 kb the catalogue is almost entirely insertions. This asymmetry has two contributions that our design cannot separate: large deletions face stronger purifying selection than insertions of comparable size, a pattern reported in other mammalian pangenomes (Li et al., 2023; Collins and Talkowski, 2025),and reference-based short-read calling systematically underascertains large insertions because the inserted sequence is absent from the linear reference (Ebler et al., 2022).The second is precisely the bias a pangenome removes, and the large insertions recovered here are the clearest example of variation that this approach makes accessible and a conventional workflow would not.

Structural variant density and gene content did not correlate, with all correlations explaining at most 12 % and typically under 2 % of the variance (*R*^2^ = 0.0002 to 0.123). This contradicts the expectation that purifying selection depletes SVs from gene-rich regions (Sudmant et al., 2015; Collins and Talkowski, 2025).The most likely explanation is ascertainment: because a pangenome captures a broader and less curated spectrum of variation than reference-based calling, including many transposable-element-derived insertions of neutral or modest effect, any negative correlation present in a high-confidence functional SV set is diluted here (Crysnanto et al., 2021; Liu, 2025).

The same SV set recovered the four population clusters as clearly as the SNP set (Groningen White Headed 0.98, Dutch Belted 1.00 at *K* = 4), indicating that structural variants can serve as population-genetic markers in their own right wherever a pangenome makes them accessible to short-read genotyping, rather than only as a complement to SNPs.

Ten structural variants were fixed within a single breed and located in annotated genes (Table 4).With five to seven animals per breed, presence in every animal of one breed is by itself weak evidence of breed-specificity, since a common variant can be carried by all sampled animals by chance. Testing each SV against the genome-wide *V*_*ST*_ distribution and against permuted breed labels showed that all ten are among the most breed-differentiated SVs in the panel rather than common variants that happened to pass the frequency filter. The clearest genotype-phenotype link is a 174 bp deletion in *KIT* on BTA6, present in all Dutch Belted individuals and absent elsewhere. *KIT* encodes a receptor tyrosine kinase governing melanocyte migration and survival, and structural variation at this locus underlies white-spotting and belting phenotypes across mammals (Giuffra et al., 1999; Andersson, 2020).The primary driver of the belt in cattle is a non-coding copy number variant upstream of *TWIST2* on BTA3 (Awasthi Mishra et al., 2017; Rothammer et al., 2018),and our windowed *F*_*ST*_ analysis recovers that locus independently; BTA3 is by far the most differentiated chromosome in the Dutch Belted comparison against Holstein Friesian, contributing 27 of 91 windows in the top 0.1 %, and the two windows immediately flanking the variant both exceed that threshold (*F*_*ST*_ = 0.658 and 0.728). The two windows spanning the variant itself fall just below the threshold and carry roughly a third as many segregating sites as their neighbours, which is informative rather than contradictory: short-read SNP genotyping loses both sensitivity and site density inside a tandem-repeat expansion, so a reference-based scan localises the region while going partially blind at the causal site. Since belt width and completeness vary considerably among belted animals and are not fully explained by *TWIST2* (Hogers et al., 2026),and since quantitative variation in *KIT* pathway activity modulates the spatial extent of melanoblast colonisation in a dose-dependent manner (Yoshida et al., 2001; Mort et al., 2016), this deletion is a plausible modifier. Recent pangenome work showing that SVs at *KIT* underlie head depigmentation in white-headed taurine breeds supports the same broader role (Sorin et al., 2026).

The Groningen White Headed carried the most breed-specific SVs, mirroring its disproportionate contribution to breed-private non-reference sequence (Table S1) and reinforcing the picture of elevated structural divergence. The largest is a 1376 bp insertion in *CPLANE1* on BTA20, fixed in the breed and present in one Dutch Friesian Red and one Meuse-Rhine-Yssel animal. *CPLANE1* and *AHI1*, which also carries a breed-specific insertion, both function in primary cilium formation, but neither these nor the variants in *DENND1B* and *AK8* can currently be linked to a breed phenotype. The *DENND1B* pair warrants caution rather than interpretation: the 66 bp insertion and 66 bp deletion, separated by 907 bp and of identical length and nearidentical sequence, has the signature of a pseudo-balanced indel pair arising from alignment ambiguity in repetitive sequence rather than of two independent mutations (D’Costa and Simpson, 2023; Ding et al., 2025).

However, our results also indicate that pangenomes are not without limitations. First, the graph represented the belted CNV as a single large insertion rather than as parallel, countable paths (Figure S1), so k-mer genotypers such as PanGenie, had no local structure against which to resolve copy number. Reference-guided builders such as Minigraph-Cactus are optimised for stability and standard structural variants and are known to struggle with multi-copy loci (Liao et al., 2023), whereas all-to-all induction methods such as PGGB (Garrison et al., 2024) may yield the topology required. Construction was not the only problem, however, our Dutch Belted assembly collapsed the animal’s four CNV copies into three, and a genotyper can only resolve copy numbers represented among the input haplotypes. Accurate pangenome-based CNV genotyping therefore requires both an appropriate graph construction method and input assemblies spanning the copy-number range present in the target population. Second, PanGenie requires non-overlapping records, so the nested bubble hierarchy must be collapsed to top-level sites; where the reference path traverses a snarl ambiguously, vg deconstruct emits no top-level record at all, and restricting the panel to top-level sites then discards that variation despite its presence in the graph. This removed 12.0 % of records overall and close to half of BTA6, including a contiguous block spanning 7.7 Mb to 64.2 Mb. The regions affected are the structurally complex ones a pangenome is best placed to represent, so the same complexity that motivates the approach can render parts of it inaccessible. Thirdly, of the 486 genotype comparisons at the nine Groningen White Headed marker positions, 471 (96.9 %) agreed between graph-based and linear-reference calling, but the 15 discordances were strongly directional, with 14 under-calling the alternate allele in the graph and 13 involving Groningen White Headed animals. At 20 044 910 bp (BTA20) the linear reference recovered the alternate allele in all seven Groningen White Headed individuals while the pan-genome called every one homozygous reference, excluding both low population frequency and ONT error as explanations and locating the failure in graph genotyping itself. That this occurs in the most divergent breed in the panel is a caution against assuming that graph-based genotyping is uniformly superior for short variants. In addition the candidate variants identified here require functional follow-up, beginning with targeted validation of the fixed breed-specific calls and extending to expression and phenotypic analysis, before they can inform selection.

### Towards pangenome-informed breeding and conservation

Taken together, our findings define a practical opportunity. Roughly 157 Mb of breed-private sequence and tens of thousands of structural variants in this panel alone are simply not addressable by a linear reference, and the variants that emerge when they are made addressable are not trivial: a deletion in a coat-colour master regulator, fixed in the breed defined by its coat pattern and the most breed-differentiated SV in that breed. Such a variant would not appear on a conventional SNP array, and structural variants of this kind are not captured by the SNP and small-indel calling on which most short-read cattle studies rely. That such a candidate emerges from eight breeds and 54 animals suggests that part of the functional variation underlying breed-defining phenotypes is not directly genotyped by the SNP arrays and short-variant pipelines commonly used for trait mapping in cattle, and is captured only indirectly where it happens to be in strong linkage disequilibrium with assayed SNPs. Graph-genotyped structural variants proved reliable enough to recover the same population structure as the SNP set, indicating that they behave as informative markers rather than noise. Whether they add value beyond SNPs remains to be tested, for example through pangenome-informed imputation into within-breed association mapping and genomic prediction: genomic selection operates largely within breeds, where the extensive linkage disequilibrium in cattle means that many SVs will be tagged by nearby SNPs and explain little additional variance, and translating SVs into a robust genotyping array poses technical challenges of its own.

For conservation the argument is more immediate, because the populations concerned may not persist. Three of these breeds retain fewer than 350 female breeding animals, and Dutch traditional breed populations declined by up to 95 % between 2020 and 2026 (Centrum voor Genetische Bronnen, 2020, 2026).Producing and storing a dedicated reference assembly for each small, closely related population would be both impractical and largely redundant, whereas one graph captures the shared backbone once and retains what distinguishes each breed. Two results here bear directly on how future sequencing efforts should be coordinated. The first is that assembly number drove the raw breed-private totals, so a panel with one assembly per breed systematically underestimates shared variation and cannot distinguish a genuinely private large insertion from an unsampled one; adding haplotypes per breed can be more prudent than adding breeds. The second is that divergence and within-breed diversity can point in opposite directions, as they do in the Groningen White Headed, so prioritisation criteria based on distinctiveness alone will misjudge which populations are closest to losing the variation they hold.

However, realising this will require improvements on three fronts. Assemblies built from higher-accuracy chemistry, R10.4.1 ONT or PacBio HiFi, would reduce the spurious short variation that inflates non-reference totals and should yield fewer unresolvable snarls. Graph construction that represents multi-copy loci as countable paths, and genotypers able to operate directly on nested bubble structures rather than requiring a flattened top-level panel, would recover the 12 % of records lost during panel preparation and make copy-number states genotypable.

## Conclusion

This study presents the first Dutch cattle pangenome, capturing 193.4 Mb of non-reference sequence from 12 *de novo* assemblies of seven traditional breeds. The majority of that sequence is breed-private, and increasingly so with variant size; variants *≥*1 kb are almost entirely lineage-specific, a pattern that persists after controlling for the number of assemblies per breed. Graph-based genotyping of a short-read cohort recovered SNPs, indels and structural variants in a single pass, and all three variant classes resolved the same four genetic clusters. This confirmed the marked divergence of the Groningen White Headed alongside its reduced within-breed diversity, and yielded 55 865 structural variants, including a *KIT* deletion fixed in the Dutch Belted that was the most breed-differentiated SV in that breed. Based on our findings from using a pangenome for conservation genetics in small populations, we can make some suggestions regarding future studies of this type. First, assembly number rather than biology drove the raw breed-private totals, so additional haplotypes per breed can be more informative than additional breeds. Second, divergence and within-breed diversity can point in opposite directions, as in the Groningen White Headed, so prioritisation on distinctiveness alone will misjudge which populations are closest to losing what they hold. Third, graph topology and the flattening required by current genotypers still limit access to complex loci; higher-accuracy assemblies, construction methods that represent multi-copy loci as countable paths, and functional validation of the candidate variants are the next steps. The resource presented here is Dutch, but the design applies wherever small, closely related populations must be characterised on a budget that cannot support a reference assembly for each.

## Data availability

Genomic data of the traditional Dutch cattle, including the *de novo* assemblies, the short- and long-reads, and the pangenome, are available at the European Nucleotide Archive under project accession ERP206176. Short-read sequencing data of the Holstein Friesian are available under project PRJEB90816 at the European Nucleotide Archive. The genome assembly workflow is available at https://codeberg.org/Rensco/Bovine-Assembly-Pipeline under the European Union Public Licence v1.2. Scripts used for read processing, pangenome construction, genotyping, and downstream analyses are available at https://codeberg.org/Rensco/Dutch_cattle_pangenome under the MIT License, and will be archived on Zenodo with a DOI upon acceptance.

## Author contributions statement

R.H. wrote the manuscript and analysed the data. A.B. and R.C. conceived the study. A.B., C.D.B., M.B, and R.C. led the project. B.D and M.G. performed the lab experiments and data storage A.R. raised the funding. All authors read and approved the final manuscript.

## Acknowledgments

We thank all members of the OPTIBOV consortium for their invaluable contributions to sample collection and sequencing. This work was supported by the EU-Africa LEAP-Agri (OPTIBOV project, LEAP-Agri-326), the European Union’s Horizon 2020 research and innovation program (grant No. 727715). We acknowledge the Dutch Belted Breeders’ Association (Vereniging lakenvelder Runderen), Groninger White headed (Blaarkop Stichting), Meuse-Rhine-Yssel Study (MRY Studieverenigingen Zuid en Oost), Deep Red Cattle (Vereniging Het Brandrode Rund), and Dutch Friesian (Fries Hollands Rundvee Stamboek). We would also like to acknowledge the Centre for Genetic Resources, the Netherlands (CGN) of Wageningen University and Research for access to sperm samples for genomic sequencing, specially Henk Sulkers and Agnes de Wit.

## Shared windowed *F*_*ST*_ peaks on BTA20

Cross-examining the windowed *F*_*ST*_ coordinates identified 12 distinct BTA20 windows shared by at least two breeds (Table S5), the most conserved lying at 42.32 Mb to 42.37 Mb and surfacing in four populations. The Groningen White Headed and Dutch Belted shared no window coordinates with any other breed, consistent with their distinct positions in the *F*_*ST*_, PCA, and ADMIXTURE analyses. Against the Groningen White Headed selection signatures reported by Gonzalez-Prendes et al. (2022), our cohort recovered 21 elevated windows within the BTA5 region (10.0 Mb to 19.7 Mb), eight within the proximal BTA20 cluster, and one within the distal cluster, with no overlap in the BTA4 or BTA25 intervals.

## Supplementary figures Supplementary tables

**Figure S1** The belted CNV is encoded as a single insertion node rather than as countable tandem copies. Graph topology at the BTA3 locus harbouring the copy number variant (CNV) underlying the belted phenotype. The CNV is represented as a single large non-reference insertion node (indicated) rather than as a series of parallel bubbles or looping paths, meaning the copy-number architecture of the locus is not explicitly encoded in the graph topology.

**Figure S2** The Groningen White Headed is the most differentiated breed, the red breeds the least. Pairwise *F*_*ST*_ heatmap showing genetic distances between the seven traditional Dutch breeds and the Holstein Friesian outgroup.

**Figure S3** A shared BTA20 signal separates six of seven traditional breeds from Holstein Friesian. Windowed pairwise *F*_*ST*_ (50 kb windows, 25 kb step) between each of the seven traditional Dutch breeds and the Holstein Friesian, calculated using the Weir and Cockerham estimator. Panels show (A) Deep Red, (B) Dutch Belted, (C) Dutch Friesian, (D) Dutch Friesian Red, (E) Groningen White Headed, (F) Improved Red, and (G) Meuse-Rhine-Yssel, each compared against Holstein Friesian. Points are coloured by chromosome (*Bos taurus* autosomes, BTA1–BTA29); the horizontal dashed line marks the top 0.1 % empirical threshold of windowed *F*_*ST*_ values for each comparison.

**Figure S4** Variance explained declines steeply through PC4 in all three variant classes. Variance explained by the first ten principal components for SNP-, indel-, and structural-variant-based PCA.

**Figure S5** Ancestry components separate the Groningen White Headed first, then the red, Friesian, and Dutch Belted groups. ADMIXTURE ancestry proportions for the eight Dutch cattle populations at *K* = 2 to *K* = 4, estimated separately from SNPs, indels and structural variants. Each vertical bar represents one individual and its ancestry proportions, which sum to one; individuals are grouped by breed (columns) and results are shown for each variant type and value of *K* (rows).

**Table S1.** Breed-private non-reference sequence stratified by node length, using all 12 *de novo* assemblies and the public Holstein Friesian genome assembly. Values represent the total sequence (Mb) in non-reference graph nodes not traversed by any assembly of another breed, partitioned by node length stratum. *n* gives the number of assemblies contributing to each breed.

| Breed | *n* | ≤5 bp | 6 bp to 49 bp | 50 bp to 199 bp | 200 bp to 999 bp | ≥1 kb | Total |
|---|---|---|---|---|---|---|---|
| Meuse-Rhine-Yssel | 4 | 11.52 | 1.52 | 2.15 | 11.46 | 36.58 | 63.23 |
| Groningen White Headed | 3 | 5.68 | 1.19 | 1.54 | 9.37 | 35.88 | 53.66 |
| Dutch Friesian Red | 1 | 1.45 | 0.31 | 0.52 | 3.21 | 10.35 | 15.84 |
| Dutch Belted | 1 | 3.73 | 0.20 | 0.11 | 0.43 | 1.96 | 6.43 |
| Improved Red | 1 | 2.43 | 0.17 | 0.16 | 0.75 | 2.42 | 5.93 |
| Deep Red | 1 | 2.95 | 0.19 | 0.09 | 0.31 | 1.25 | 4.79 |
| Holstein Friesian | 1 | 0.62 | 0.11 | 0.11 | 0.74 | 2.42 | 4.00 |
| Dutch Friesian | 1 | 1.70 | 0.14 | 0.08 | 0.28 | 1.15 | 3.35 |

**Table S2.**
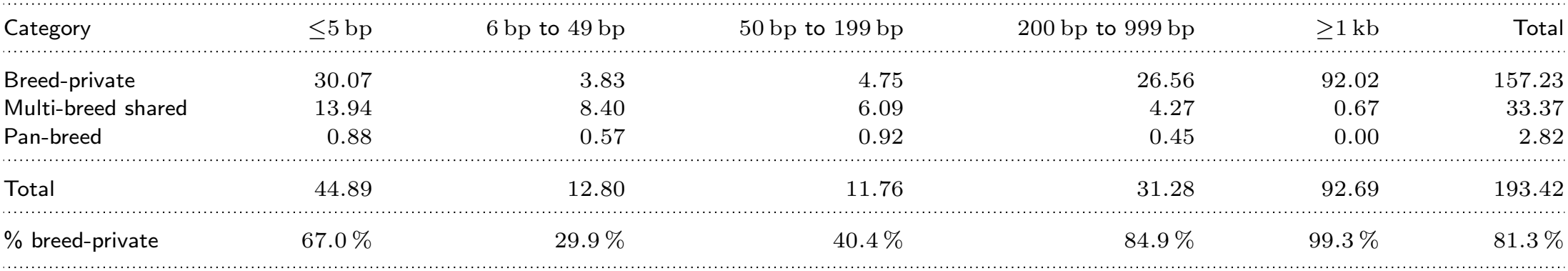
Genome-wide non-reference sequence (Mb) by sharing category and node length stratum. Breed-private: traversed by assemblies of one breed only. Multi-breed shared: traversed by assemblies of more than one but not all breeds. Pan-breed: traversed by at least one assembly of every breed. Percentages indicate the breed-private fraction of total sequence at each stratum.

**Table S3.** Breed-private non-reference sequence stratified by node length, after downsampling to a single assembly per breed. Breeds represented by several assemblies accumulate breed-private sequence partly through assembly number alone, which this table corrects.

| Breed | ≤5 bp | 6 bp to 49 bp | 50 bp to 199 bp | 200 bp to 999 bp | ≥1 kb | Total (Mb) |
|---|---|---|---|---|---|---|
| Dutch Friesian Red | 1.80 | 0.56 | 0.75 | 3.49 | 10.48 | 17.08 |
| Dutch Belted | 4.07 | 0.40 | 0.25 | 0.50 | 1.97 | 7.19 |
| Improved Red | 2.78 | 0.35 | 0.33 | 0.90 | 2.43 | 6.79 |
| Deep Red | 3.29 | 0.37 | 0.22 | 0.38 | 1.26 | 5.52 |
| Holstein Friesian | 0.94 | 0.23 | 0.23 | 0.91 | 2.50 | 4.81 |
| Groningen White Headed | 2.07 | 0.37 | 0.32 | 0.53 | 1.47 | 4.76 |
| Meuse-Rhine-Yssel | 2.40 | 0.30 | 0.21 | 0.38 | 1.20 | 4.49 |
| Dutch Friesian | 1.97 | 0.27 | 0.19 | 0.37 | 1.16 | 3.96 |

**Table S4.** Presence of SNP positions strongly associated with the GWH (taken from Table 2 of (Gonzalez-Prendes et al., 2022)) within *de novo* assembled genomes in the pangenome and short-read samples of all included Dutch cattle breeds. DBE: Dutch Belted, DF: Dutch Friesian, DFR: Dutch Friesian Red, DR: Deep Red, GWH: Groningen White Headed.

| **Position on BTA20 (bp)** | **Affected gene** | **Assemblies with variant** | **Short-read samples with variant** |
|---|---|---|---|
| 17 974 182 | *ZSWIM6* | 1/3 GWH, 1/1 DFR | 7/7 GWH, 1/7 DBE |
| 18 330 187 | *NDUFAF2* | 1/3 GWH, 1/1 DFR | 5/7 GWH, 1/7 DFR |
| 20 044 595 | *PDE4D* | 2/3 GWH | 5/7 GWH |
| 20 044 910 | *PDE4D* | 2/3 GWH | 0/7 GWH, 1/7 DF |
| 20 278 747 | *PDE4D* | 2/3 GWH | 7/7 GWH |
| 20 314 517 | *PDE4D* | 2/3 GWH | 7/7 GWH |
| 20 524 538 | *RAB3C* | 1/3 GWH | 7/7 GWH |
| 20 542 032 | *RAB3C* | 2/3 GWH | 7/7 GWH |
| 20 598 559 | *RAB3C* | 2/3 GWH | 7/7 GWH |

**Table S5.**
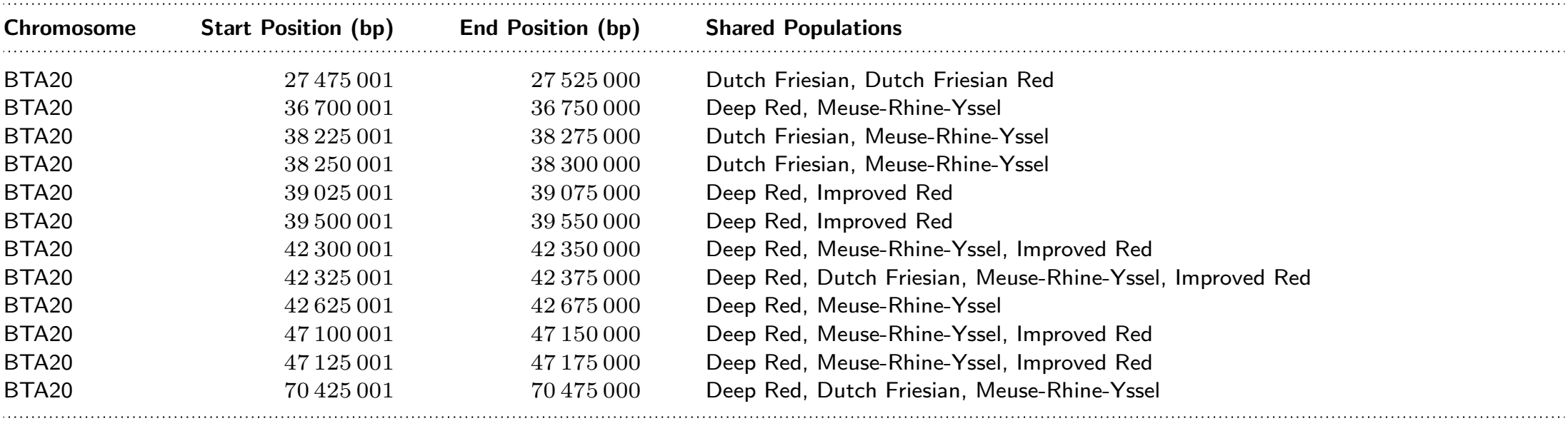
Summary of overlapping windowed *F*_*ST*_ peaks on BTA20 among traditional Dutch cattle breeds.

